# Unique nigral and cortical pathways implicated by epigenomic and transcriptional analyses in a rotenone rat model of Parkinson’s disease

**DOI:** 10.1101/2024.12.17.628873

**Authors:** Maria Tsalenchuk, Kyle Farmer, Sandra Castro, Abigail Scheirer, Yuqian Ye, J. Timothy Greenamyre, Emily M. Rocha, Sarah J. Marzi

## Abstract

Pesticide exposure is increasingly recognised as a potential environmental factor contributing to the onset of idiopathic Parkinson’s disease, yet the molecular mechanisms underlying this connection remain unclear. This study aims to explore how pesticide exposure disrupts key brain regions involved in Parkinson’s disease pathology by reshaping gene regulatory landscapes. Using the well-established rotenone rat model of the disease, we performed H3K27ac ChIP-sequencing to profile active regulatory elements in the substantia nigra and cortex. In this model, rotenone distributes uniformly throughout the brain, and the degree of complex I inhibition is equivalent in cortical and substantia nigra neurons. Despite the uniformity of complex I inhibition, we identified widespread epigenomic differences, with brain region specific acetylation patterns associated with rotenone exposure. We showed consistent changes in transcriptomic activity by RNA-sequencing. Our results indicate there is a strong immune response to rotenone localised to the substantia nigra and highlight an enrichment of immune-related motifs in this brain region, suggesting that the immune response is at least partially driven by gene regulatory mechanisms. We also noted an increase in C1q complement pathway activity in the substantia nigra. In contrast, we identified widespread dysregulation of synaptic function at the gene regulatory level in the cortex of these same rats. Our results highlight a role for gene regulatory mechanisms potentially mediating the effects of pesticide exposure, driving region-specific functional responses in the brain that may contribute to the pathology and selective vulnerability that characterise Parkinson’s disease.

## Introduction

Parkinson’s disease (PD) is the fastest growing neurological disorder globally (Dorsey et al., 2018). It is marked by progressive neurodegeneration of dopaminergic neurons in the substantia nigra pars compacta (SN) leading to the loss of dopamine in the striatum, neuroinflammatory activation of glial cells and aggregation of phosphorylated α-synuclein (Kalia and Lang, 2015). While the precise causes of PD remain unclear, a combination of genetic and environmental factors is believed to drive its onset and progression. Environmental exposures, including pesticides, have been increasingly recognised as important contributors to idiopathic PD (Freire and Koifman, 2012). However, the molecular mechanisms through which these environmental agents contribute to neurodegeneration are not yet fully understood.

Pesticide exposure, particularly mitochondrial complex I inhibitors like rotenone, induce a PD-like pathology in animal models of the disease, recapitulating many of the motor and non-motor symptoms observed in PD patients (Betarbet et al., 2000). In adult rats, rotenone administration reproduces several pathogenic pathways, including oxidative stress, α-synuclein (α-syn) phosphorylation and aggregation in surviving SN dopaminergic neurons and in the gastrointestinal tract, lysosomal and proteasomal dysfunction and nigral iron accumulation (Cannon et al., 2009; Rocha et al., 2020; Cannon et al., 2009). Additionally, rotenone induces systemic changes affecting both the brain and peripheral tissues, which is significant as PD is increasingly recognised as a multi-system disorder. Studies have shown that rotenone exposure can lead to gastrointestinal issues and cardiac dysfunction (Zhan et al., 2020; Johnson et al., 2018), mirroring non-motor symptoms observed in PD patients (Skjærbæk et al., 2021; Kincl et al., 2024).

Gene regulation and chromatin dynamics are increasingly recognised as central mechanisms in the development of neurodegenerative diseases (Marzi et al., 2018). Epigenetic modifications, such as histone acetylation, play a role in regulating gene expression and maintaining cellular identity. Epigenetic investigations in PD remain scarce, with only one study delving into the role of H3K27ac in the cortex of PD patients (Toker et al., 2021). Understanding these regulatory alterations in response to environmental insults is crucial for identifying molecular drivers of PD (Tsalenchuk et al., 2023).

In this study, we explored how rotenone exposure alters the epigenetic and transcriptomic signatures of two key brain regions involved in PD pathology: the SN and the cortex. This approach not only seeks to uncover potential vulnerabilities in the SN compared to the cortex but also aims to provide a comprehensive understanding of regional PD pathology. As the disease progresses, PD patients often experience cognitive impairment and dementia, alongside gastrointestinal and motor impairments (Hanagasi et al., 2017). These cognitive changes are thought to be associated with cortical pathology (Yu et al., 2019; Braak et al., 2006; Irwin et al., 2017). Here, we used H3K27ac chromatin immunoprecipitation (ChIP)-seq and RNA sequencing (RNA-seq) to profile gene regulatory variation in the SN and cortex of rats exposed to rotenone compared to vehicle controls. Our results reveal widespread epigenetic and transcriptomic changes, highlighting distinct functional responses across different brain regions. This study represents the first comprehensive assessment of gene regulatory alterations in the rotenone rat model.

## Results

Aged (8-10 months) male Lewis rats received daily systemic exposure to rotenone for 21 days (ChIP-seq; N = 9, RNA-seq; N = 20). At the end of the dosing regimen, rats have extensive PD pathology in the brain. To confirm rotenone caused a nigrostriatal lesion, tyrosine hydroxylase-positive terminals were assessed in the striatum (**Supplementary Figure 1**). Subsequently, we dissected the cortex and SN to characterise the transcriptomic and epigenetic patterns caused by rotenone. To profile the histone acetylation landscape, we conducted ChIP-seq for H3K27ac, which marks active promoters and enhancers across the genome. To link our epigenetic data to variation in gene expression levels, we performed RNA-seq to profile mRNA transcripts in a separate set of animals (**Figure 1a**).

**Figure 1:**
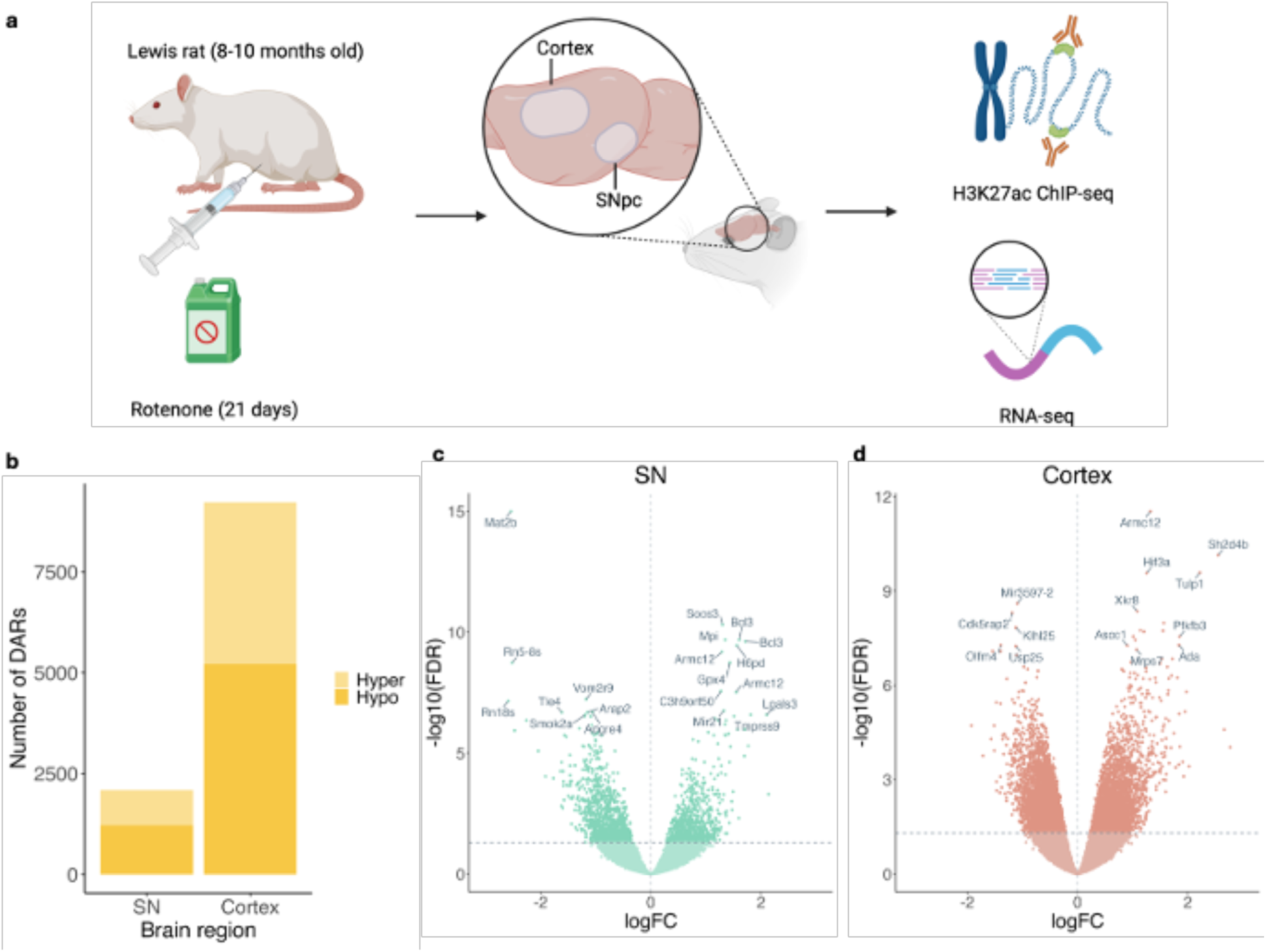
Epigenomic profiling of rotenone exposed rat cortex and SN reveals widespread changes in H3K27 acetylation. **a)** Experimental workflow: Male Lewis rats were exposed to 2.8 mg/kg rotenone or vehicle intraperitoneally daily for 21 days. 24-hrs following the final injection, the cortex and SN regions were dissected from each brain (n = 5 for rotenone-treated, n = 4 for vehicle-treated, per brain region). ChIP-seq targeting H3K27ac was performed on these samples, complemented by RNA-seq. **b)** Stacked bar plot showing the number of differentially acetylated regions (DARs) identified in the cortex and SN. **c)** Volcano plot showing DARs in the SN. **d)** Volcano plot showing DARs in the cortex.

After stringent quality control and pre-processing, we obtained high-quality H3K27ac ChIP-seq data from the SN and cortex of nine rats (rotenone n = 5, vehicle n = 4) and RNA-seq data from 10 SN and 8 cortex samples each from rats treated with rotenone or vehicle (**Figure 1a**). On average, we obtained 32,790,067 sequencing reads per ChIP-seq sample (SD = 21,518,441) with no significant difference in read depth between the rotenone-exposed groups and controls (p-value = 0.54). In total we called 38,281 peaks in the SN and 80,763 in the cortex. We observed widespread differences in active promoters and enhancers across both brain regions in response to rotenone, accompanied by corresponding changes in gene expression. In the SN, 874 peaks showed increased acetylation and 1216 showed reduced acetylation upon rotenone exposure (**Supplementary Table 1 and 2**), with significant hypoacetylated peaks being more prevalent (3.18% vs 2.28%, 3.33 x 10^-14^, exact binomial test). In the cortex, 4016 peaks were hyperacetylated and 5213 were hypoacetylated upon rotenone exposure (**Supplementary Table 3 and 4**), with 11.4% of the peaks showing differential acetylation and a higher frequency of hypoacetylation (p-value = 2.2 x 10^-16^, exact binomial test). Of note, two of the top hyperacetylated peaks in the SN were annotated to *BCL3:* a promoter peak (logFC = 1.61, p = 2.18 x 10⁻¹⁴) and an intronic peak (logFC = 1.71, p = 3.20 x 10⁻¹⁴) (**Supplementary Figure 2**). *BCL3*, a gene involved in immune signalling, has been implicated in late-onset Alzheimer’s disease and is linked to hyperinflammation in microglia (Wightman et al., 2021). Notably, *BCL3* was also one of the most overexpressed genes in the SN (logFC = 5.33, p < 4.42 x 10^13^) (**Supplementary Table 5**). Conversely, the top hypoacetylated peak in the SN was annotated to *MAT2B*, a gene involved in coenzyme metabolic processes, which has been found to be downregulated in patients with Lewy Body dementia (Santpere et al., 2017). One of the strongest hyperacetylated signatures in the cortex was observed for *HIF3A*, a gene typically upregulated in response to hypoxia in the brain (Heidbreder et al., 2003), while *OLFM4*, a gene involved in regulating synaptic function and inflammation (Xu et al., 2023) was found to be hypoacetylated.

To reveal dysregulation in biological pathways upon exposure to rotenone, we investigated functional enrichments amongst differentially acetylated regions and differentially expressed genes using pathway enrichment analysis (**Figure 2**). We identified several consistently altered pathways across both brain regions, including upregulation of GO terms “cellular response to peptide hormone stimulus” (SN; p = 3.11 x 10^-7^, cortex; p = 4.25 x 10^-15^) and “actin filament organization” (SN; p = 3.08 x 10^-7^, cortex; p = 1.62 x 10^-14^), which may indicate a response to mitochondrial dysfunction and associated damage to the axon and cytoskeleton (Zhang et al., 2023). However, not all pathways were consistent between brain regions. In the cortex, other enriched terms in the hyperacetylated peaks included those associated with epithelial cell migration (epithelial cell migration; p = 5.69 x 10^-15^, epithelium migration; p = 8.14 x 10^-15^) potentially representing activation of survival pathways in response to rotenone (**Figure 2c**). In contrast, in the SN, pathways associated with oxidative stress (response to hydrogen peroxide; p = 1.55 x 10^-9^, response to reactive oxygen species; p = 4.57 x 10^-9^) were among the most enriched in the hyperacetylated peaks (**Figure 2a**). The pathways also diverged in the hypoacetylated peaks. The SN showed downregulation of pathways associated with Wnt signalling (Wnt signaling pathway; p = 6.24 x 10^-6^, cell-cell signaling by Wnt; p = 6.91 x 10^-6^) (**Figure 2b**), while the cortex showed downregulation of pathways associated with regulation of neurogenesis (regulation of neurogenesis; p = 4.07 x 10^-16^) (**Figure 2d**). The pathway enrichment analysis appears to show a stronger correlation between ChIP and RNA data in the SN compared to the cortex. For an overall assessment of the concordance between ChIP-seq and RNA-seq, we correlated logFC values between promoter-associated DARs and corresponding gene expression (**Supplementary Figure 3**). We observed a significant correlation between ChIP-seq and RNA-seq data, with a stronger association in the SN samples (R = 0.57) compared to the cortex samples (R = 0.42). This indicates a general concordance between changes in H3K27 acetylation and gene expression.

**Figure 2:**
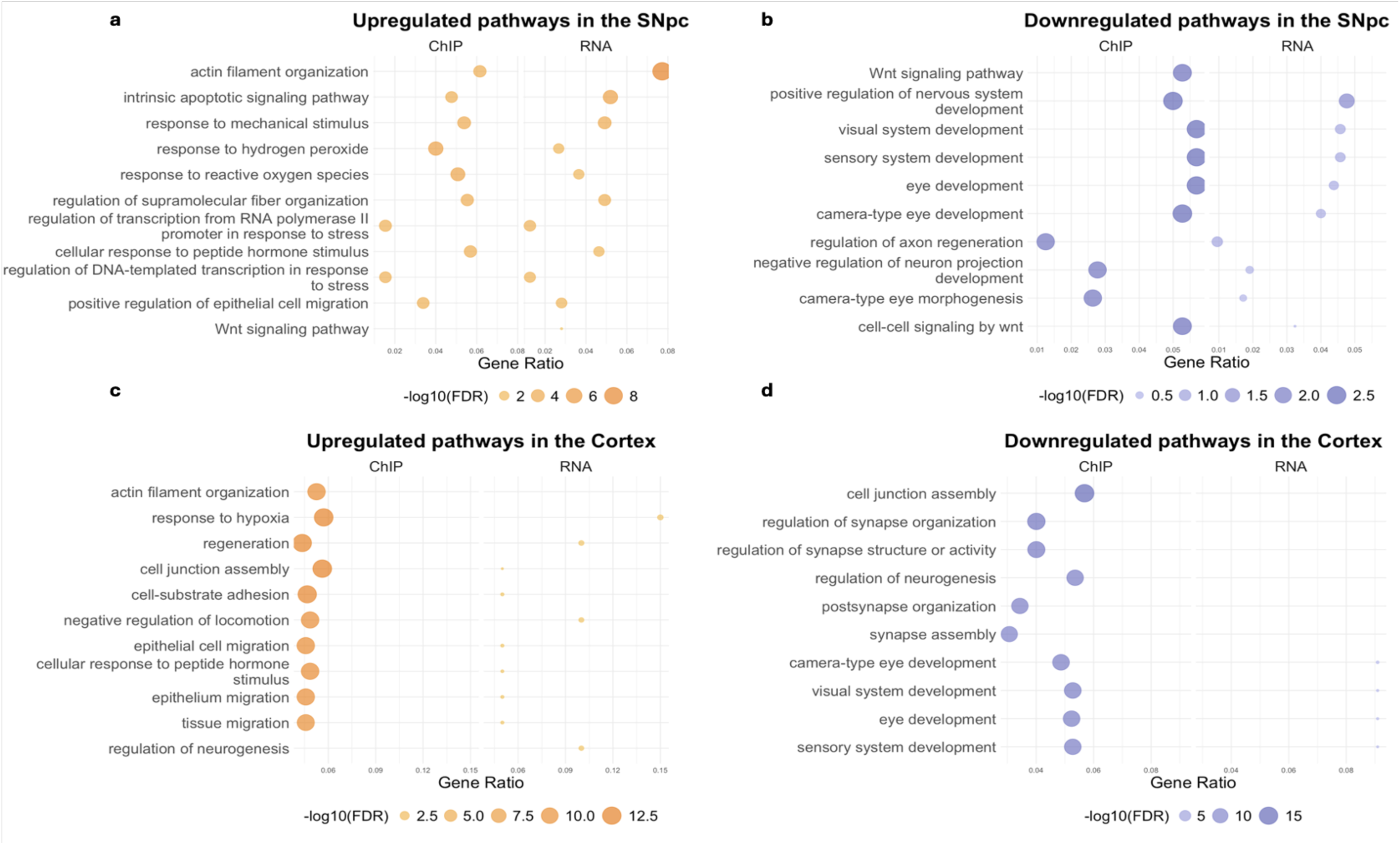
Pathway enrichment analysis reveals dysregulation of pathways linked to stress response and Wnt signalling in the SN, and hypoxia and synapse organisation in the cortex. Dot plots illustrating **a)** upregulated pathways in the SN, **b)** downregulated pathways in the SN, **c)** upregulated pathways in the cortex, and **d)** downregulated pathways in the cortex. Notably, there appears to be a stronger correlation between ChIP and RNA data in the SN.

### Transcriptomic analysis with RNA-seq supports gene regulatory responses identified from histone acetylation signatures

To complement the histone acetylation signatures seen in both regions exposed to rotenone with transcriptomic activity, we performed low coverage RNA-seq (average number of reads = 953,647, SD = 516,771) on ten samples from the SN per condition and eight samples from the cortex per condition. We identified widespread changes in the SN transcriptome, where 791 genes were upregulated and 652 downregulated (FDR range = 6.68 × 10−13 - 4.99 × 10−2, log2FC range = −3.62 - 7.65), following exposure to rotenone (**Figure 3a**). In the cortex, 21 genes were found to be upregulated, and 13 genes were downregulated (FDR range = 6.15× 10−5 - 4.38 × 10−2, log2FC range = −2.63 – 1.92) (**Figure 3b**). To investigate whether genes were showing concerted expression changes across gene regulatory networks, we used weighted gene co-expression network analysis (WGCNA) to classify and analyse gene modules. These modules were then tested for differences in expression between the rotenone and control groups. In the cortex, only one gene module showed significant differential expression, whereas in the SN, seven modules were downregulated, and one was upregulated (**Figure 3c,d**). Pathway enrichment analysis revealed that the downregulated module in the cortex was associated with a disturbance in cellular respiration as evidenced by enriched terms such as oxygen transport (p = 1.42 x 10^-8^), carbon dioxide transport (p = 4.65 x 10^-9^) and gas transport (p = 6.86 x 10^-8^). The decrease in peroxidase activity (p = 9.08 x 10^-5^) and oxidoreductase activity (p = 1.01 x 10^-4^), also points towards alterations in oxidative stress response mechanisms (**Figure 3e**). In contrast, the most significantly downregulated module in the SN was associated with cellular respiration, with all terms referring to mitochondrial processes, for example, mitochondrial respiratory chain complex assembly (p = 7.49 x 10^-20^) and NADH dehydrogenase complex assembly (p = 7.12 x 10^-18^) (**Figure 3f**). Meanwhile, a module upregulated in the SN pertained to protein synthesis, including terms like ribonucleoprotein complex binding (p = 1.84 x 10^-13^), ribosome biogenesis (FDR = 1.01 x 10^-9^) and regulation of translation (FDR = 3.56 x 10^-11^) (**Figure 3g**). This is an intriguing observation as previous research has indicated a decrease in RNA translation in response to rotenone exposure and in fibroblasts of *LRRK2* G2019S PD patients (Deshpande et al., 2020). Therefore, our data may suggest a compensatory RNA translation mechanism in response to rotenone.

**Figure 3:**
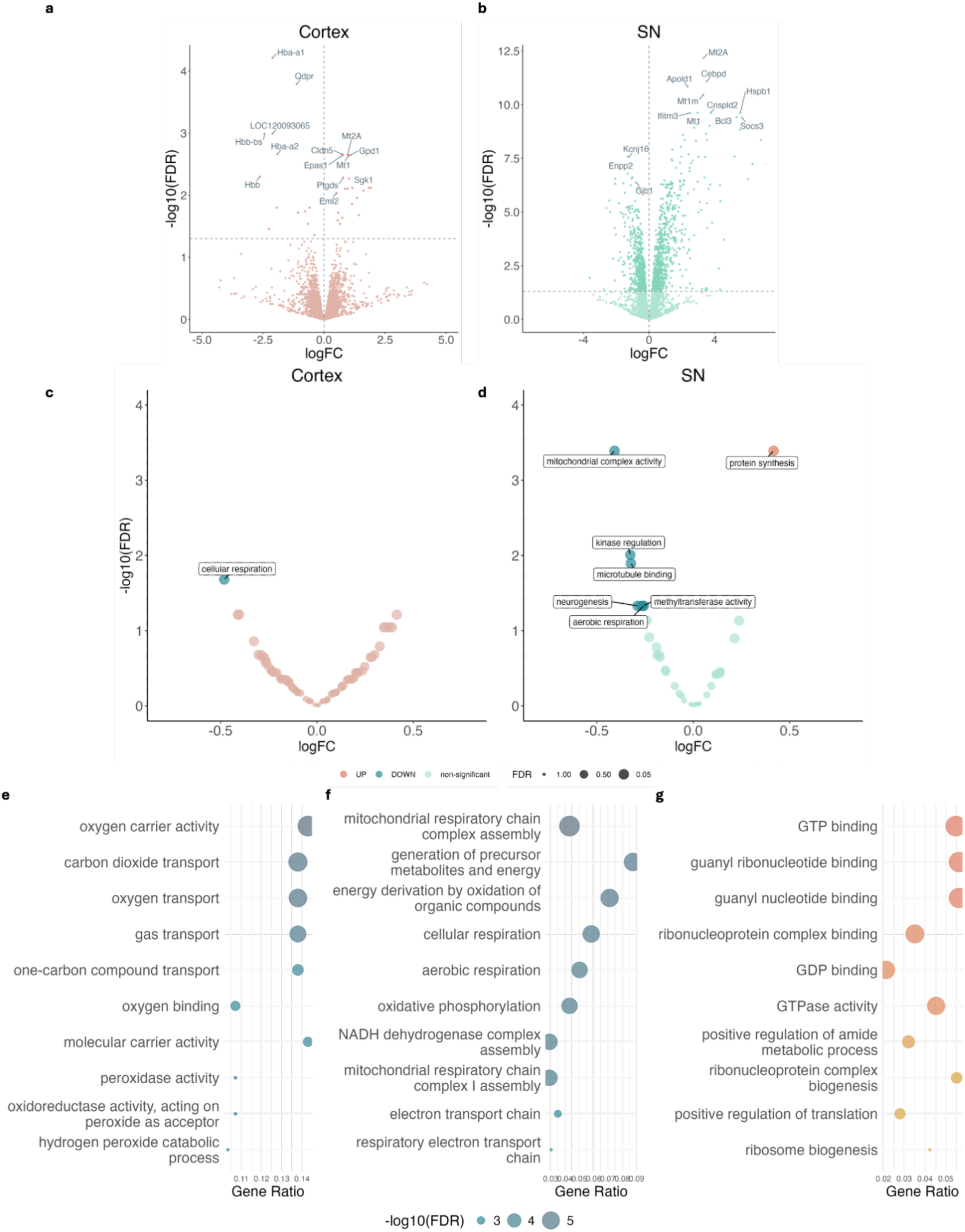
Gene co-expression network analysis reveals the upregulation of protein synthesis pathways and the downregulation of mitochondrial function-related gene modules in the SN. Volcano plot of differentially expressed genes in **a)** the cortex and **b)** the SN. Volcano plot of differentially expressed gene modules in **c)** the cortex and **d)** the SN. **e)** Dotplot depicting enriched pathways in the downregulated module in the cortex. **f)** Dotplot depicting enriched pathways in the most significantly downregulated module in the SN. **g)** Dotplot depicting enriched pathways in the upregulated module in the SN.

### Differential acetylation patterns reveal unique rotenone-associated immune-related changes in the SN

To identify pathways unique to each brain region, we performed an overlap of all the peaks from the cortex and SN. A total of 24,878 peaks were shared between the two brain regions (**Figure 3a,b**). Among these shared peaks, the cortex exhibited 1,788 significantly hyperacetylated regions and 2,224 significantly hypoacetylated regions (FDR range = 5.96 × 10−9 - 4.99 × 10−2, log2FC range = −2.53 – 1.41). In contrast, the SN showed 643 hyperacetylated peaks and 890 hypoacetylated peaks (FDR range = 2.72 × 10−11 - 4.99 × 10−2, log2FC range = −1.63 – 2.72). Of the significant peaks (FDR < 0.05) in both SN and cortex, the direction of effect was the same for 80% of peaks. It should be noted there are more overall reads in the cortex than the SN, so there is more power to detect differences, which may account for the disparity in number of DARs between brain regions. Pathway analysis across consistently altered peaks in both brain regions re-confirmed upregulation of the pathway “cellular response to peptide hormone stimulus” (cortex; p = 8.69 x 10^-13^, SN; p = 4.33 x 10^-8^).

When looking at the peaks unique to each brain region, 5,205 peaks were found to be unique to the SN. Of these, 136 were hyperacetylated and 250 were hypoacetylated (**Figure 3a**). Intriguingly, there were several hyperacetylated pathways that were uniquely enriched in the SN in response to rotenone exposure (**Figure 3b**). These pathways were associated with the immune response, including chemotaxis (p = 5.77 x 10^-6^), activated T cell proliferation (p = 2.74 x 10^-5^), leukocyte apoptotic process (p = 6.00 x 10^-5^) and myeloid leukocyte activation (p = 7.29 x 10^-5^). These findings suggest a heightened immune and cellular activation response associated with rotenone exposure, which seems to be unique to the SN. These findings are consistent with published reports demonstrating rotenone induced SN inflammation (Rocha et al., 2022). T-cell activation is particularly interesting, given that in PD, autoimmune responses by T cells to α-syn are strongest in the earlier stages of disease and attenuate 10 years after diagnosis (Lindestam Arlehamn et al., 2020). Chemotaxis also implies an immune response wherein microglia migrate towards the site of inflammation in response to cytokines (Fan et al., 2017). Furthermore, we identified a peak annotated to *CXCL11* among the top hyperacetylated peaks unique to the SN. *CXCL11* is a cytokine, which is known for its chemotactic properties for activated T cells (Burns et al., 2006).

To further our understanding of the upstream regulators of the rotenone response, we performed transcription factor motif enrichment analysis using HOMER on significantly hyper- and hypoacetylated peaks SN (**Figure 3c, Supplementary Table 6 and 7**). Strikingly, and in agreement with the observed pathway enrichments, regions within the SN exhibiting increased acetylation were found to be enriched with immune-related transcription factor motifs. This included PU.1, the IRF family, RUNX1, the STAT family, GATA3, and IL-21. This result suggests heightened activity of transcription factors that regulate immune function in the SN of rotenone-exposed rats. Using a gene list of transcription factors involved in immune function and differentiation (Liu et al., 2022), we identified which motifs were immune related and examined whether their corresponding genes showed differential expression or acetylation in the SN (**Figure 3d**). Concordant changes in both gene expression and binding sites could indicate biological relevance, while the expression signature may help distinguish the binding patterns of these transcription factors, despite their overlapping motifs. We found that the majority of transcription factors (including *Fos*, *Irf8, Stat3* and *Maff*) showed both increased expression and elevated H3K27ac levels. *Irf8* is particularly interesting in the context of PD: Knock-down of *IRF8* has been shown to alleviate neuroinflammation and behavioural deficits in a MPTP mouse model of PD. This suggests that IRF8 may be a critical mediator of inflammatory processes driving neurodegeneration in PD (Ma et al., 2024).

Expanding on the immune response observed in the SN following rotenone exposure, we identified the complement system as upregulated in both epigenomic and transcriptomic analyses (**Figure 4**). The complement system, a crucial part of the innate immune response, is responsible for recognizing and clearing dying cells (Bajic et al., 2015). It has also been implicated in abnormal synaptic pruning in brain region-specific models of AD, where inhibition of C1q rescues synaptic loss. In this study, the SN of exposed rats showed cumulative hyperacetylation of 56 genes previously reported to be involved in the complement system (**Figure 4a**) (Carpanini et al., 2021). Of note the A-, B- and C-chains of complement component q1 (C1q) of the classical complement pathway are upregulated both at the level of gene expression (**Figure 4b**) and acetylation in the SN but not cortex of rotenone-exposed rats (**Figure 4c, Supplementary Table 8**).

**Figure 4:**
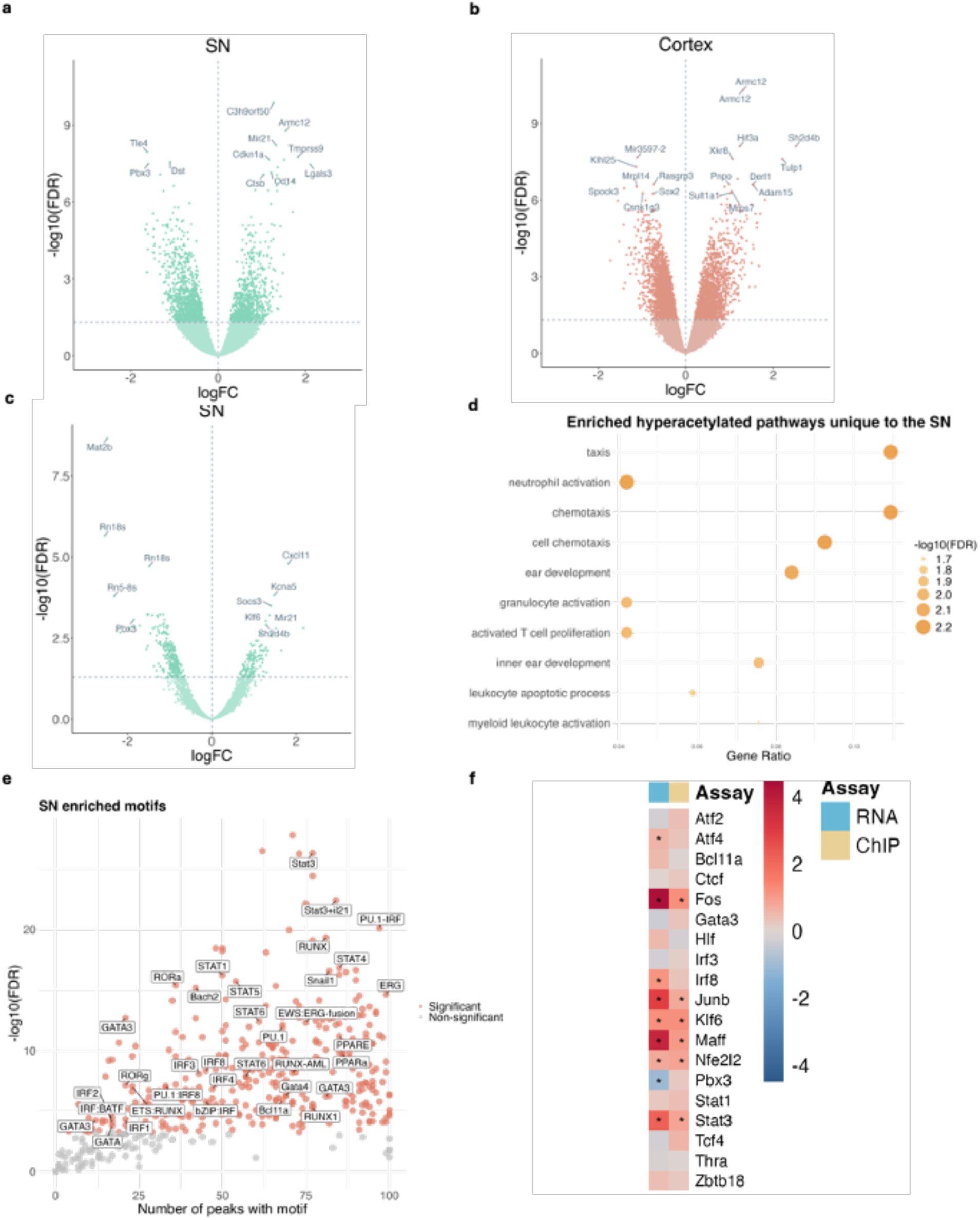
Immune genes show upregulation at both the transcriptional and epigenetic levels in the SN following exposure to rotenone. **a)** Volcano plot of peaks shared between cortex and SN, with values specific to the SN shown. **b)** Volcano plot of peaks shared between cortex, with values specific to the cortex shown. **c)** Volcano plot of H3K27ac peaks unique to the SN. **d)** Pathway enrichment of hyperacetylated peaks unique to the SN. **e)** Motif analysis shows enrichment of immune motifs (labelled) in hyperacetylated peaks in the SN. **f)** Heatmap showing log fold change for corresponding H3K27 acetylation and gene expression of immune-related transcription factors. Stars indicate significant log fold change (FDR < 0.05).

**Figure 5:**
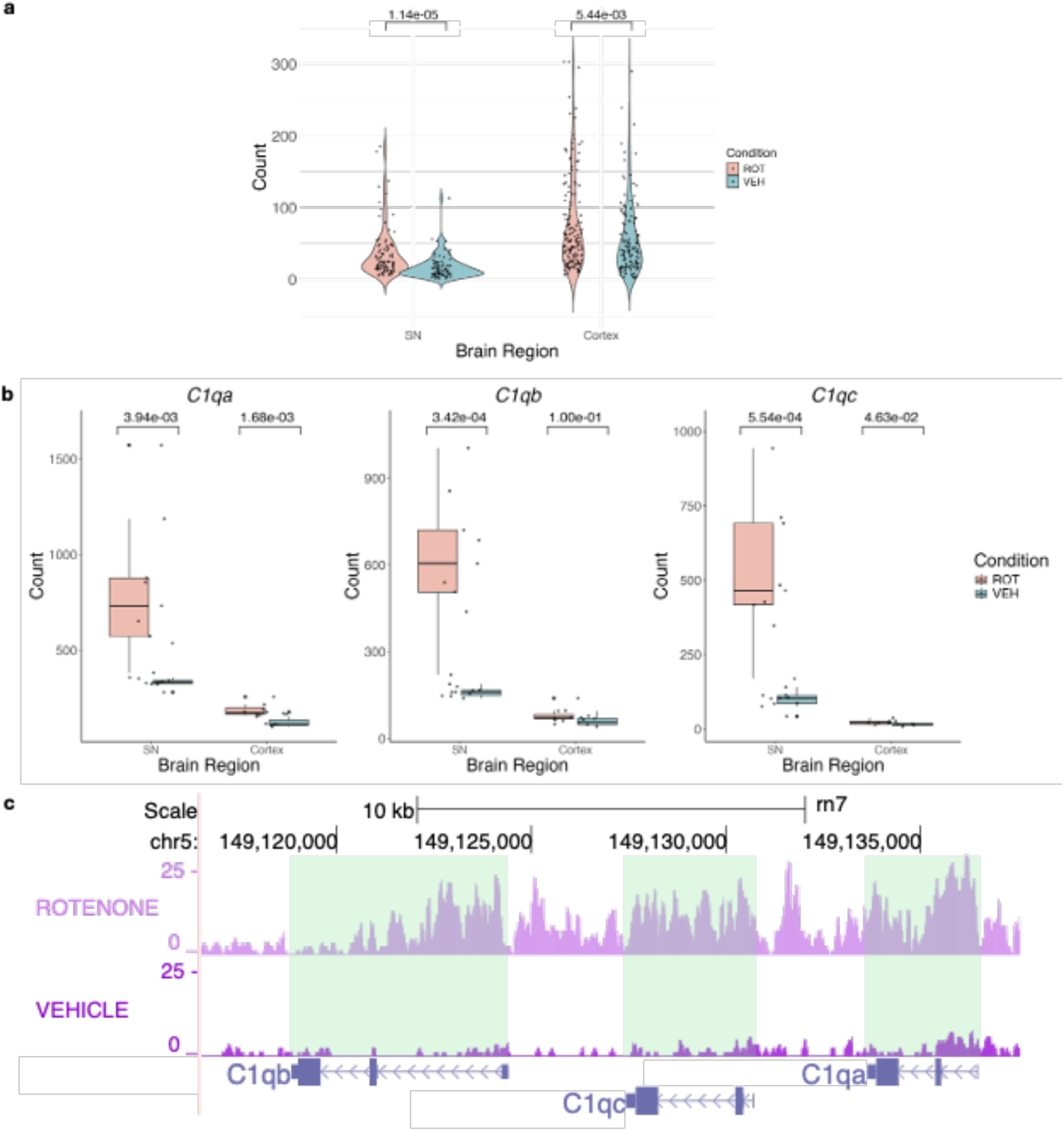
Upregulation of the complement system genes in the SN following rotenone exposure. **a)** Violin plot showing increased acetylation of complement system genes in the SN and cortex of rotenone-treated samples. **b)** Boxplot displaying elevated gene expression of the A-, B- and C-chains of C1q in the SN of the rotenone-treated group. **c)** Genome browser track showing higher H3K27ac signals in C1q chain genes in rotenone SN samples compared to controls.

### Epigenetic analyses highlight synaptic dysregulation in cortex following rotenone exposure

We identified 36,296 acetylation peaks unique to the cortex, including 1,394 hyperacetylated and 1,655 hypoacetylated peaks (**Figure 6a**). Unlike in the SN, pathways enriched in cortex-specific acetylation peaks suggest dysregulation of synaptic genes following rotenone exposure. Both hyper- and hypoacetylated regions are associated with processes such as synapse assembly (hyper; p = 2.61 x 10⁻^10^), regulation of synapse organisation (hypo; p = 1.90 x 10⁻^11^), and postsynapse organisation (hypo; p = 5.98 x 10^⁻11^) (**Figure 6b**).

**Figure 6:**
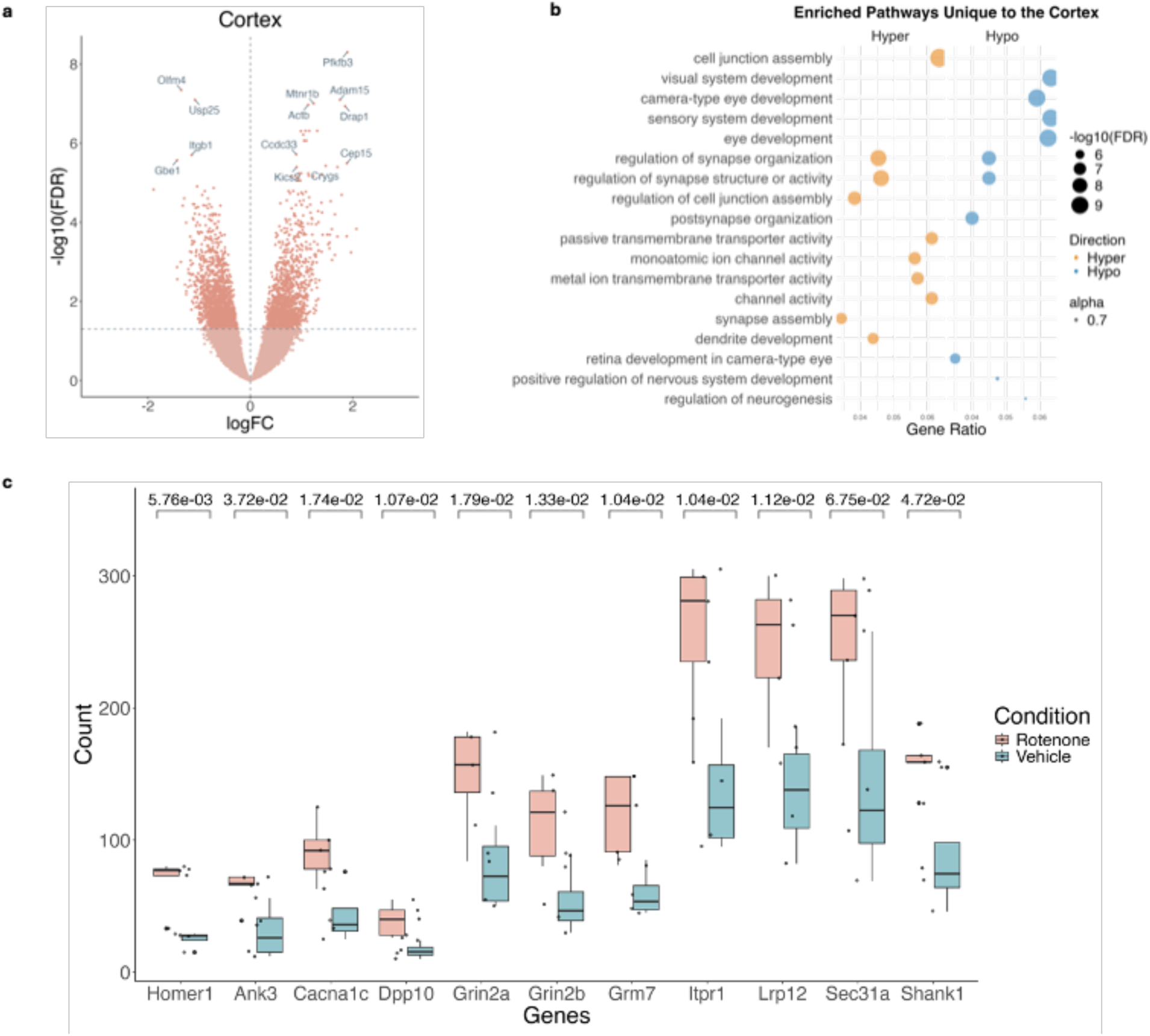
Dysregulation of synaptic pathways in the cortex upon exposure to rotenone. **a)** Volcano plot of H3K27ac peaks unique to the cortex. **b)** Pathway enrichment of hyper- and hypo- acetylated peaks unique to the cortex. **c)** Box plots showing hyperacetylation of Homer1 and its known interactors in the cortex, showing the most significant peak for each gene.

Among these hyperacetylated genes are *HOMER1, GRIN2B,* and *SHANK1*, which are known to interact (Stillman et al., 2022; Yoon et al., 2021). HOMER1 directly binds to GRIN2B, and its hyperacetylation may enhance this interaction, potentially leading to dysregulation of NMDA receptor activity, a pathway implicated in excitotoxicity. Moreover, HOMER1 scaffolds with SHANK1 at excitatory synapses, where it plays a critical role in maintaining dendritic spine integrity. The hyperacetylation of HOMER1 may stabilise these synaptic complexes in response to rotenone-induced stress, serving as a compensatory mechanism.

However, given HOMER1’s broader role in synaptic protein networks, particularly through its interactions with proteins containing the Homer1-binding PPXXF motif, such as ankyrin-G (Yoon et al., 2021), hyperacetylation could excessively stabilise these complexes. This may drive overactive excitatory signalling, further contributing to excitotoxicity, a well-known mechanism linked to PD progression (Iovino et al., 2020). Supporting this, we also observe hyperacetylation in genes encoding postsynaptic proteins that contain the PPXXF motif and are known to bind Homer1 (**Figure 6c**).

Importantly, several of these genes have been previously implicated in PD. HOMER1 has been linked to psychotic symptoms in PD through genetic mutations (De Luca et al., 2009), while GRIN2B polymorphisms have been associated with impulse control behaviours in PD (Zainal Abidin et al., 2015). Additionally, other genes that interact with Homer1 have been associated with PD, including *CACNA1C* (Wang et al., 2016), *GRIN2A* (Yamada-Fowler et al., 2014), *SEC31A* (Antoniou et al., 2022), *LRP12*, and *SHANK1* (Kaiser et al., 2023).

## Discussion

Rotenone is a pesticide, also readily used to control nuisance fish populations, and serves as a well-established agent for inducing PD-like symptoms and pathology in rat models. To induce a Parkinsonian phenotype in rats, the toxin is administered systemically (intra-peritoneal cavity) and inhibits mitochondrial complex I uniformly throughout the brain. Strikingly, rotenone causes selective dopaminergic neurodegeneration in the SN despite reaching other dopaminergic neuron populations (Rocha et al., 2022). The underlying mechanism responsible for this selective vulnerability is not clear, therefore we sought to better understand rotenone-induced epigenetic and transcriptomic changes in the SN. Identifying the source of this selective vulnerability will provide insight into the underlying causes of PD and may identify new therapeutic avenues. In this study, we quantified levels of histone acetylation (H3K27ac) and gene expression in the SN and cortex of rats exposed to rotenone, revealing widespread variation in both assays.

Consistent with known mechanisms implicated in rotenone toxicity, our findings underscore oxidative stress and mitochondrial impairment as core elements of its neurotoxic effects (Rocha et al., 2022; Testa et al., 2005). Oxidative stress and mitochondrial dysfunction can exacerbate each other, contributing to a cascade that impairs neuronal survival and function (Guo et al., 2013). In line with this, differential H3K27 acetylation analysis identified genes implicated in oxidative stress, as upregulated in both the SN and cortex. Mitochondrial dysfunction is strongly reflected in our transcriptomic data, which shows downregulation of gene modules linked to mitochondrial function and cellular respiration, as identified through WGCNA. The gene expression changes we observed in both the cortex and SN are consistent with previous findings, further supporting the downregulation of mitochondrial pathways. For example, rotenone exposure in induced pluripotent stem cell-derived dopaminergic neurons led to a reduction in oxidative phosphorylation-related genes, as shown by single-cell RNA-seq (Fernandes et al., 2020). Mitochondrial dysfunction and oxidative stress have also been noted in extra-nigral areas such as the striatum and cortex (Thomas-Broome and Castorina, 2022). Additionally, another study found upregulation of oxidative stress genes across various tissues, including the brain (Heinz et al., 2017). In our data, oxidative stress pathways were especially enriched in the SN, suggesting a potential role in dopaminergic neuron loss in this region. This is notable as it underscores a differential gene-level response to oxidative stress between the cortex and SN.

Of note, epigenetically upregulated genes in the SN showed enrichment in immune response pathways, while this response was not observed in the cortex. This is consistent with previous evidence demonstrating heightened immune response in the SN, as indicated by highly reactive microglia (Stefanova, 2022). Rotenone has been shown to trigger a midbrain-restricted inflammatory response, with heightened expression of glial markers observed only in the nigra, and not in extra-nigral regions in rodent models (Rocha et al., 2022; Thomas-Broome and Castorina, 2022; Van Laar et al., 2023). The immune response in PD has been recognized since the 1920s, initially observed through an increase in the number of reactive microglia (Schonhoff et al., 2020). Post-mortem brain samples of PD patients have revealed increased levels of the pro-inflammatory cytokine TNF-α and the T cell-associated chemokines CXCL12 in the SN, preceding the loss of dopaminergic neurons (Shimoji et al., 2009), a pattern that aligns with our observations of H3K27ac in rotenone-exposed rats. In a rotenone mouse model, specific microglial activation in the SN was observed in response to rotenone, although the cortex, striatum, and globus pallidus did not exhibit a corresponding increase in immune activity (Rocha et al., 2022). Our data suggests that the immune response may be driven by the variety of immune related transcription factors identified by the motif analysis, including the STAT family, IRF8 and PU.1, although additional validation is warranted to confirm this hypothesis. We also observed a striking, nigra-specific upregulation of both acetylation and expression of *C1QA*, *C1QB*, and *C1QC*, key components of the classical complement pathway, suggesting an enhanced inflammatory response in this region.

In contrast, analysis of cortical H3K27ac data indicated alterations in synapse function. While limited research exists on the response to rotenone in the cortex, a study demonstrated that primary cortical cells exposed to rotenone showed impaired axonogenesis (Bisbal and Sanchez, 2019). Another study using neocortical neurons treated with rotenone revealed mitochondrial dysfunction and an increase in apoptotic genes (Yap et al., 2013). Our data underscore the necessity for further investigations in the cortex, given the significant alterations we observed, particularly of synaptic pathways at the epigenetic level. Furthermore, cortical pathology is evident in PD patients, including cortical atrophy and reduced synaptic density, which progresses from the SN to cortical regions as the disease advances (Sarasso et al., 2021; Frigerio et al., 2024), mirroring observations in the cortex ChIP-seq data. Pathway analyses of altered histone acetylation following rotenone exposure suggest that rotenone-associated synaptic dysfunction may be regulated at least partially at the epigenetic level.

Our study is subject to several limitations that warrant consideration. Firstly, there is an imbalance in sequencing depth between the ChIP-seq and RNA-seq samples. While the ChIP-seq samples were sequenced at a depth of 30 million reads, the RNA-seq samples were sequenced at a depth of only 5 million reads per sample. This discrepancy in sequencing depth may have led to the underrepresentation of certain transcripts in the RNA-seq data, potentially reducing our ability to fully characterise the underlying transcriptional mechanisms associated with rotenone toxicity. In light of this, we performed primary analyses on the H3K27ac ChIP-seq data and used the low-coverage gene expression to contextualise and interpret results observed at the epigenetic level. Of note, while we detected more DARs in cortex, the ChIP-seq data had higher read coverage in the cortex compared to the SN, increasing our power to detect differentially acetylated regions and could partially explain the different number of DARs between the two brain regions. Secondly, the use of bulk tissue genomic techniques provides an averaged view of histone acetylation and gene expression across the constituent cell types under investigation. This approach may obscure potential cellular heterogeneity within the sampled tissue, limiting our ability to detect subtle changes in gene regulation that may be specific to certain cell types or subregions. Of particular relevance in this context are dopaminergic neurons and microglia, both of which occur at low proportions in the surveyed brain regions. In this context, it is of note that we are able to detect a strong SN-specific immune response in the bulk tissue samples. Additionally, the relatively small number of samples used in our study may have impacted the statistical power of our findings. While the rotenone rat model is well-established, we acknowledge that no single model can recapitulate human disease. Advances in machine learning for cross-species translation of genomic results, are promising in this regard (Marzi et al., 2023). Finally, while our study focused on H3K27ac as a marker of active enhancers, it is important to acknowledge that other epigenetic mechanisms, such as DNA methylation or histone methylation, could also play a role in mediating the effects of rotenone on gene expression and neuronal function. Future studies incorporating a broader range of epigenetic markers may provide a more comprehensive understanding of the molecular mechanisms underlying rotenone-induced neurotoxicity.

In summary, our study provides compelling evidence linking alterations in histone acetylation and gene expression to rotenone exposure. Our findings indicate that brain region-specific immune responses and synaptic dysregulation are key factors contributing to the neurotoxic effects of rotenone in the brain, and we show that this is potentially mediated by epigenetic mechanisms.

## Methods

### Rotenone rat generation

All experiments using animals were approved by the Institutional Animal Care and Use Committee of the University of Pittsburgh. Male Lewis rats aged 8-10 months received either rotenone at a dose of 2.8 mg/kg or a vehicle solution (Miglyol) administered intraperitoneally once daily for 21 days (Rocha et al., 2020). For ChIP-seq experiments, five rats were exposed to rotenone and four rats were in the control group for both brain regions. In each treatment group for RNA-seq experiments of the SN, there were 10 rats, while for the cortex, there were eight rats per treatment group. 24 hours after the final rotenone injection, rats were perfused using a saline-buffered solution. The cortex and SN were microdissected and immediately placed on dry ice.

### Nuclei isolation for ChIP-seq

Nuclei isolation from rat midbrain and cortical tissue was performed following a previously published protocol with slight modifications (Nott et al., 2021). Briefly, fresh frozen brain tissue weighing approximately 2-10 mg was homogenised using a Kimble Dounce tissue grinder in 1% formaldehyde in phosphate buffered saline (PBS). To quench the fixation, glycine was added at a final concentration of 0.125 M. The homogenate was then pelleted at 1,100 x*g* for 5 minutes at 4°C. Subsequent steps were carried out on ice or at 4°C. The homogenates were washed twice with NF1 buffer (10 mM Tris-HCl pH 7.4, 1 mM EDTA pH 8.0, 5mM MgCl_2_, 0.1 M sucrose, 0.5% (vol/vol) Triton X-100) and then incubated in 5 mL of NF1 buffer for 30 minutes. The homogenates were passed through a 70-μm cell strainer. Homogenates were underlaid with a 1.2 M sucrose cushion (1.17 M sucrose, 10 mM Tris-HCl pH 7.4, 3 mM MgCl_2_, 1 mM DTT), before centrifuging at 3,900 x *g* for 30 minutes at 4°C with the brake set to “low.” The pelleted nuclei were washed with NF1 buffer, followed by a wash with FANS buffer (PBS, 1% (wt/vol) bovine serum albumin (BSA) and 1mM EDTA pH 8.0).

### ChIP-seq library preparation

ChIP-seq was conducted based on a previously published protocol with minor modifications (Texari *et al*., 2021). Fixed nuclei were resuspended in LB3 buffer (10 mM Tris-HCl pH 7.4, 100 mM NaCl, 1 mM EDTA PH 8.0, 0.5 mM EGTA, 0.1% sodium deoxycholate, 0.5% N-Lauroylsarcosine, 1X protease inhibitor), and then transferred to Covaris microtubes with AFA fiber. All subsequent steps were carried out on ice or at 4°C. The samples were sonicated using a Covaris E220 focused-ultrasonicator (Covaris, MA) for 10 minutes (Duty: 5, PIP: 140, Cycles: 200, AMP/Vel/Dwell: 0.0). To serve as DNA input controls, 1% of the sample was stored at −20°C. For the ChIP procedure, 25 µL Protein G Dynabeads (Invitrogen, 10004D) and 3 µL H3K27ac antibody (Active Motif, 39133) were added to the samples, and rotated at 4°C overnight. The Dynabeads were subsequently washed three times with WB1 (20 mM Tris-HCl pH 7.4, 150 mM NaCl, 2 mM EDTA pH 8.0, 0.1% SDS, 1% Triton X-100), three times with WB3 (10 mM Tris-HCl pH 7.4, 250 mM LiCl, 1% Triton X-100, 1mM EDTA PH 8.0, 0.7% sodium deoxycholate), three times with TET (10 mM Tris-HCl pH 7.4, 1 mM EDTA PH 8.0, 0.2% Tween20), and finally with Te-NaCl (10 mM Tris-HCl pH 7.4, 1mM EDTA PH 8.0, 50 mM NaCl). The beads were resuspended in 25 μL of TT (10 mM Tris-HCl pH 7.4, 0.05% Tween 20). Input samples were adjusted to 25 μL with TT, and input libraries were generated in parallel with the ChIP libraries. For library preparation, the NEBNext Ultra II DNA Library Prep kit (New England BioLabs, E7645) was used for End Prep and Adapter Ligation, following the manufacturer’s instructions. Unique barcoded adapters (NextFlex, Bioo Scientific) were added. To decrosslink, libraries were incubated with RNase A (Thermo Scientific, EN0531) and Proteinase K (New England BioLabs, P8107) at 55°C for 1 hour and 65°C overnight. Libraries were then PCR amplified for 14 cycles with NEBNext Ultra II Q5® Master Mix (New England BioLabs, M0544). Gel extraction using a 10% TBE gel was performed to size-select libraries for 200-500 bp fragments. Samples were sequenced by the Imperial BRC Genomics Facility on an Illumina NextSeq2000 platform using paired-end sequencing and a read length of 75bp, targeting a depth of 30 million reads.

### RNA-seq library preparation

Samples were homogenised in 1 mL of Trizol reagent (Thermofisher Scientific, Waltham, MA) and total RNA was extracted using the Life Technologies min-RNA purification kit (Qiagen, Valencia, CA) according to the manufacturer’s protocol. Libraries were constructed using the 3’QuantSeq kit (Lexogen) with the following modifications. To enable the removal of PCR duplications, a custom oligo-dT primer was designed using a 12-bp unique molecular identifier. Samples were sequenced to a depth of 5 million, single-end reads, each on a Illumina Novaseq platform (Illumina, San Diego, CA).

### Data analysis

Initial screening for sample anomalies was performed with FastQC (version 0.11.9). Adapter sequences were trimmed using Trim Galore! (version 0.6.7). For ChIP-seq, the trimmed FASTQ files were then mapped to the rat reference genome (mRatBN7.2) using Bowtie2 (version 2.4.4), followed by the removal of duplicate reads with Picard (version 2.6.0). Peak calling was conducted using MACS2 (version 2.2.7.1) on merged samples. For normalisation, we used merged control inputs pooled from multiple animals. Specifically, reads from individual replicates were combined into a single dataset prior to peak calling, allowing us to detect regions consistently enriched across samples. The pooled input control ensured accurate background normalisation across the merged data. The parameters for calling peaks were set for paired-end BAM files, using a genome size of 2.7 billion, a shift size of 50, and all duplicate reads were retained. Finally, FeatureCounts (version 2.0.5) quantified the alignment of reads to these identified peak regions. For RNA-seq, FASTQ files were aligned, and transcripts were quantified using Salmon (version V1.10.0). Differential analysis to pinpoint epigenomic and transcriptomic differences between rotenone and control groups was performed using a quasi-likelihood F test in EdgeR (4.0.16). Normalisation factors were calculated based on sample-specific library compositions, and a quasi-likelihood model was fitted to the data using glmQLFit() and glmQLFTest() functions. Effect sizes are reported as log(fold change), which refers to the log_2_-transformed ratio of normalised counts between cases and controls. Transcripts and H3K27ac peaks with FDR-corrected p value < 0.05 were considered differentially expressed (upregulated or downregulated) or differentially acetylated (hyperacetylated or hypoacetylated) between the two experimental groups. Weighted gene co-expression network analysis (WGCNA) was performed on transcript counts in the cortex and SN respectively using the WGCNA R package (1.72.5). The H3K27ac peaks were assigned to genes using the annotatePeak() function in ChIPseeker (1.38.0) using “org.Rn.eg.db” as the annotation package and taking into account regulatory regions within 3 kb of the transcription start site. Gene ontology enrichment was performed using the enrichGO() function of the package clusterProfiler (4.10.0) for gene ontology enrichment analysis. Motif analysis was performed using HOMER, using all detectable H3K27ac peaks as background. Detailed annotated code used for analysis can be found at https://github.com/Marzi-lab/rotenone_rat_ChIP/.

## Data and code availability

FASTQ files have been deposited in the NCBI’s Gene Expression Omnibus (Edgar et al., 2002) and are accessible through GEO Series accession number GSE280519 for RNA-seq dataset and GSE280520 ChIP-seq dataset. All the data and code required to reproduce the figures in this manuscript are available in our GitHub repository: https://github.com/Marzi-lab/rotenone_rat_ChIP/.

## Contributions

SJM, MT, EMR and JTG designed the study. MT, KF, SC, and AS performed experiments. MT and YY analysed the results. MT and SJM wrote the manuscript with input from all coauthors. SJM and EMR supervised the project.

## Acknowledgements

Figure 1a was created using BioRender. ChIP-sequencing was performed by the Imperial BRC Genomics Facility.

## Supplementary figures

**Supplementary Figure 1.**
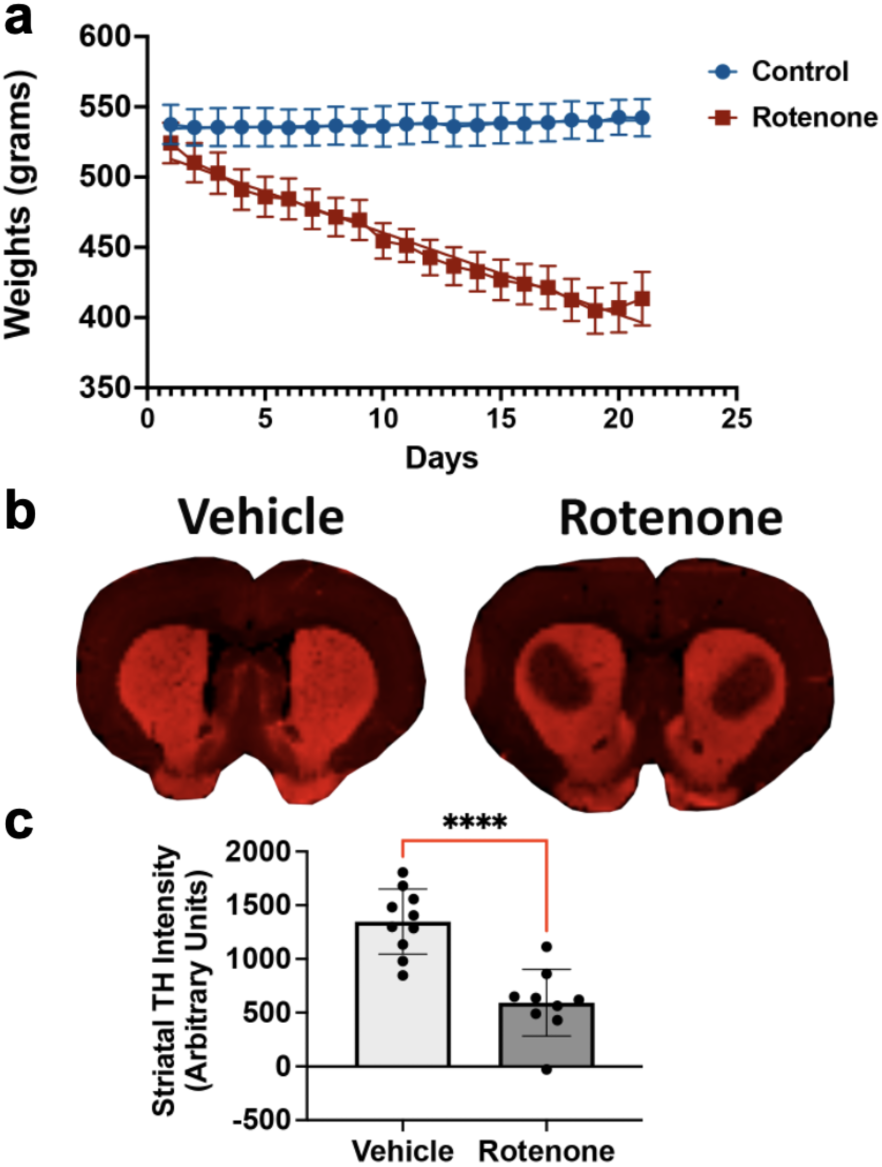
Rotenone caused a loss of striatal dopaminergic terminals. Male Lewis rats (8-10 months of age) received a single dose of vehicle (Miglyol) or rotenone (2.8 mg/kg) for 21 days. **a)** Lewis rats dosed with rotenone consistently lost weight beginning on day 3. **b-c)** Striatal dopamine terminal loss was assessed by immunohistochemistry using specific markers; tyrosine hydrolase (TH) for dopaminergic neurons. **b)** Representative photomicrographs of striatal dopaminergic terminals of rats treated with vehicle or rotenone. **c)** Quantification of striatal dopaminergic terminals loss in rats; n=10/grp. Data are analyzed using unpaired T-test, *** p < 0.0001 graphs are expressed as mean ± SEM. Symbols represent individual brains.

**Supplementary Figure 2:**
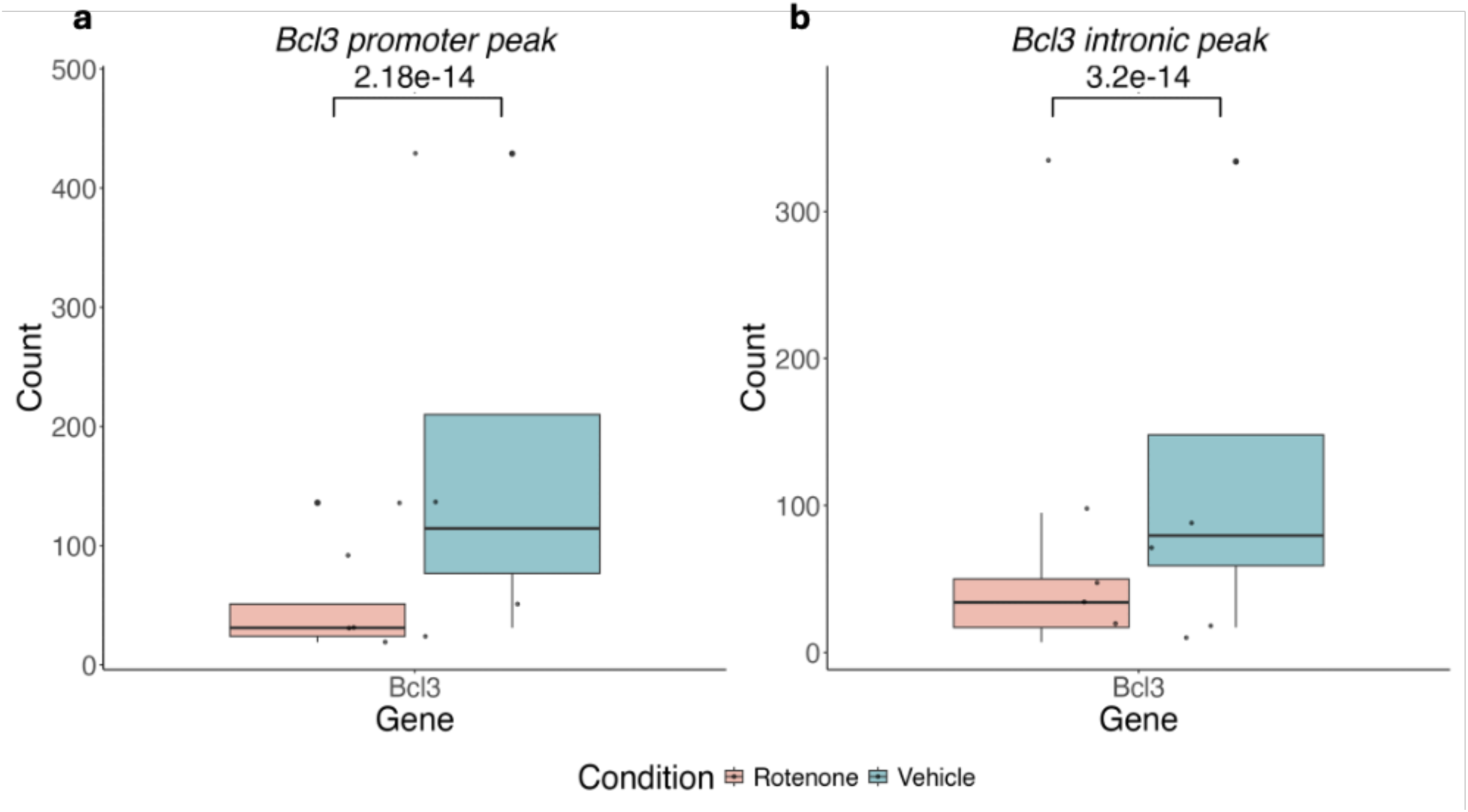
Box plots showing the hyperacetylation levels of two *Bcl3* peaks in the SN following rotenone exposure. **a)** Promoter peak. **b)** Intronic peak.

**Supplementary Figure 3:**
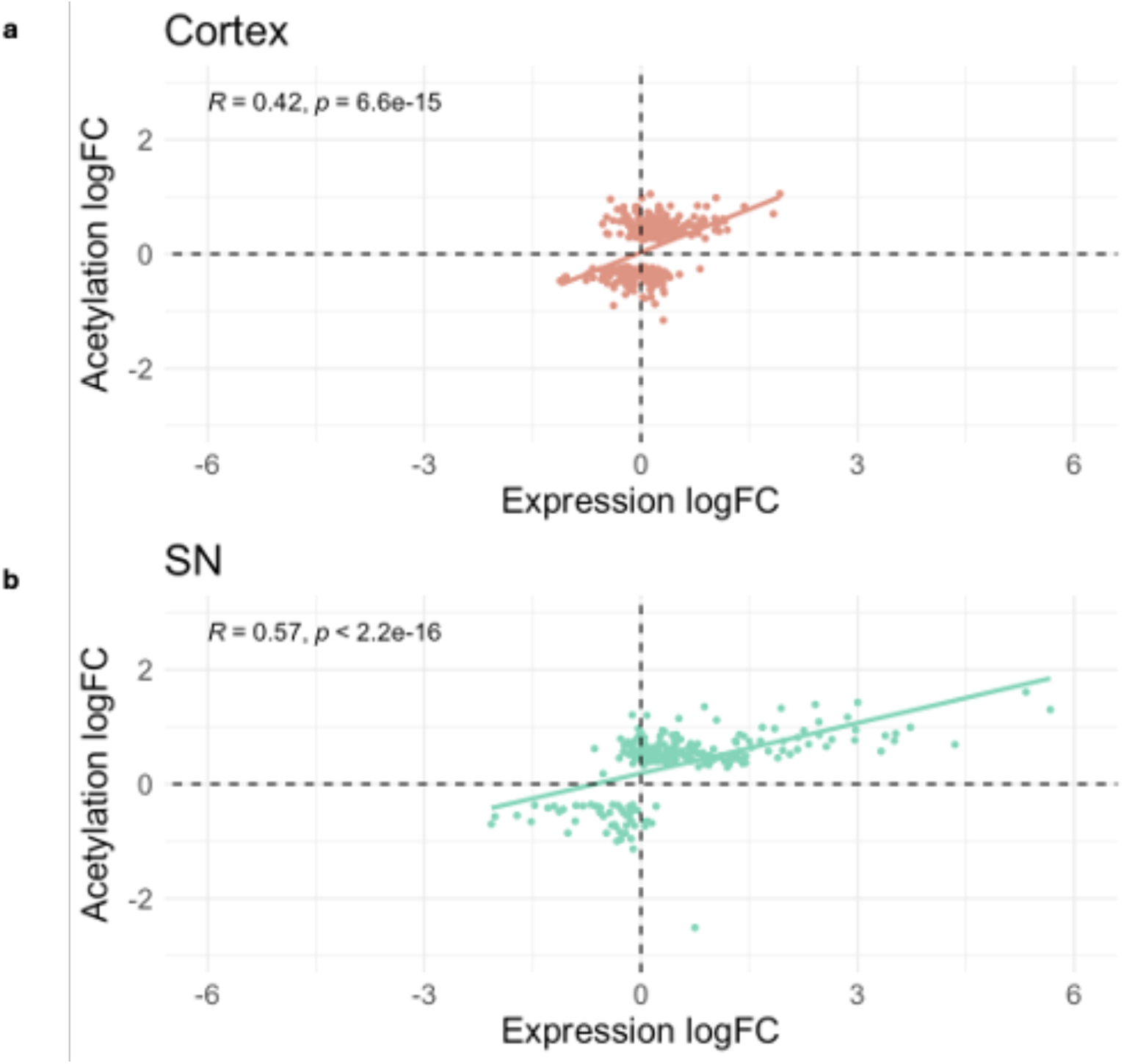
RNA-seq and ChIP-seq signals correlate across differentially acetylated promoters and their genes. Correlation of effect sizes (logFC) in acetylation and gene expression, comparing rotenone-exposed rats to vehicle controls in **a**) cortex and **b**) SN. Correlations were performed using the Pearson’s product moment correlation coefficient.

